# Homology-guided identification of a conserved motif linking the antiviral functions of IFITM3 to its oligomeric state

**DOI:** 10.1101/2020.05.14.096891

**Authors:** Kazi Rahman, Charles A. Coomer, Saliha Majdoul, Selena Ding, Sergi Padilla-Parra, Alex A. Compton

**Author notes:** These authors contributed equally (Kazi Rahman, Charles A. Coomer).

## Abstract

The interferon-inducible transmembrane (IFITM) proteins belong to the Dispanin/CD225 family and inhibit diverse virus infections. IFITM3 reduces membrane fusion between cells and virions through a poorly characterized mechanism. We identified a GxxxG motif in many CD225 proteins, including IFITM3 and proline rich transmembrane protein 2 (PRRT2). Mutation of PRRT2, a regulator of neurotransmitter release, at glycine-305 was previously linked to paroxysmal neurological disorders in humans. Here, we show that glycine-305 and the homologous site in IFITM3, glycine-95, drive protein oligomerization from within a GxxxG motif. Mutation of glycine-95 in IFITM3 disrupted its oligomerization and reduced its antiviral activity against Influenza A and HIV-1. The oligomerization-defective variant was used to reveal that IFITM3 promotes membrane rigidity in a glycine-95-dependent manner. Furthermore, a compound which counteracts virus inhibition by IFITM3, amphotericin B, prevented the IFITM3-mediated rigidification of membranes. Overall, these data suggest that IFITM3 oligomers inhibit virus-cell fusion by promoting membrane rigidity.

## Introduction

The intrinsic protection of cells from virus infection represents an early and essential aspect of antiviral innate immunity. Cytokines including interferons signal the presence of invading viruses and induce an “antiviral state” via the expression of hundreds of antiviral genes [1, 2]. This arsenal of antiviral proteins converges on many steps of the virus life cycle in order to collectively inhibit infection of cells and prevent virus spread. In addition, certain “front-line” antiviral proteins impose a constant barrier to infection because they are expressed constitutively and are further upregulated by interferons. The interferon-induced transmembrane (IFITM) proteins are the earliest acting restriction factors known, inhibiting the entry of diverse viruses into cells by restricting fusion pore formation during virus-cell membrane fusion [3-6]. Among the growing list of viruses shown to be inhibited by IFITM proteins in cell culture and in vivo are orthomyxoviruses, flaviviruses, filoviruses, alphaviruses, and coronaviruses [3]. IFITM3 is a potent inhibitor of Influenza A virus (IAV) infection in cell culture and in vivo, and consequently, it is the most studied member of the IFITM family [7-9]. While the precise mechanism by which IFITM3 reduces virus-cell fusion remains unresolved, evidence suggests that it does so by altering the properties of lipid membranes.

Two models have been proposed to explain how IFITM3 inhibits virus fusion. In the first, IFITM3 plays an indirect role by interacting with VAMP-associated protein A (VAPA) and inhibiting lipid transport between the endoplasmic reticulum and endosomes, resulting in an accumulation of endosomal cholesterol [10]. High cholesterol content may inhibit the fusion of virus-containing vesicles with the limiting membrane of the late endosome, restricting virus release into host cell cytoplasm. While inhibition of cholesterol trafficking has been shown to inhibit virus entry [11], this model of IFITM3 function has been subsequently supported by one study [12] but challenged by several others [13-16]. The second and currently favored model posits that IFITM3 directly inhibits fusion by locally reducing membrane fluidity (i.e. increasing membrane rigidity or order) [14, 17] and by inducing positive membrane curvature [17]. Membrane order and curvature influence one another [18-20] and, together, they regulate numerous membrane fusion processes [21]. Consistent with the notion that IFITM3 induces local membrane order and curvature to disfavor virus-cell fusion, IFITM3 contains a juxtamembrane amphipathic helix which is essential for antiviral activity [22]. Amphipathic helices have been identified in many eukaryotic, prokaryotic, and viral proteins and are well known for their ability to bind and bend membranes [23, 24]. Another piece of evidence that supports the membrane deformation model of fusion inhibition is that IFITM3 incorporated into enveloped viruses also impairs virion fusion with target cells [25-32]. Despite this progress, a complete mechanistic view of IFITM3 is lacking because studies describing the impact of IFITM3 on membranes have not included mutants lacking antiviral function.

It was previously reported that IFITM3 forms clusters on virus-containing vesicles [33] and that IFITM3 oligomerization promotes restriction of IAV [34]. However, the determinants initially purported to mediate oligomerization (phenylalanines at residues 75 and 78 of the CD225 domain [34]) were later shown to be unnecessary when oligomerization was measured using a FRET-based approach in living cells [35]. Therefore, the oligomerization of IFITM3 appears to be influenced by unknown determinants and its importance to antiviral function is not established.

In the current study, we set out to identify a loss-of-function mutation in IFITM3 suitable for mechanistic studies by using a homology-guided approach. The *IFITM* genes (*IFITM1, IFITM2, IFITM3, IFITM5*, and *IFITM10* in humans) are members of an extended gene family known as the Dispanin/CD225 family (hereafter referred to as CD225 proteins) [36, 37]. Members of this group are characterized by the presence of a CD225 domain but the functions of most remain unknown. However, one member is the subject of numerous studies because it is linked to neurological disorders. Mutations in proline-rich transmembrane protein 2 (PRRT2) result in conditions of involuntary movement, such as paroxysmal kinesigenic dyskinesia, benign familial infantile seizures, and episodic ataxia [38, 39]. PRRT2 is a neuron-specific protein that is localized to pre-synaptic terminals and which inhibits synaptic vesicle fusion [40-42]. Molecular studies of disease-associated missense mutations in *PRRT2* (G305W/R) indicate that it causes loss-of-function, leading to unchecked neurotransmitter release [43-47]. Interestingly, the homologous residue in human IFITM3 is also subject to rare allelic variation in humans (G95W/R) and this mutation results in partial loss of activity against IAV infection [34]. However, the reason why this site is essential for the respective functions of PRRT2 and IFITM3 was unknown.

Here, we demonstrate that glycine-95 of human IFITM3 resides within a ^91^GxxxG^95^ motif that is highly conserved among vertebrate IFITM3 orthologs as well as PRRT2. Mutation of glycine-95 rendered IFITM3 less active against entry of IAV and vesicular stomatitis virus (VSV). Using fluorescence lifetime imaging microscopy (FLIM) and Förster resonance energy transfer (FRET) of fluorophore-tagged IFITM3, we found that the ^91^GxxxG^95^ motif mediates IFITM3 oligomerization in intact cells and that IFITM3 oligomers are mostly dimers. A mutant exhibiting loss of antiviral function (G95L) was deficient for oligomerization, indicating that IFITM3 oligomerization and restriction of virus entry are functionally associated. We leveraged this loss-of-function mutant to identify mechanistic correlates of antiviral function which are dependent upon IFITM3 oligomerization. Using a FLIM-based reporter, we found that IFITM3 increased membrane order, as previously suggested, while the G95L mutation failed to do so. These data indicate that membrane stiffening by IFITM3 oligomers is required for antiviral activity. In an effort to further probe the importance of membrane order in the antiviral mechanism, we demonstrate that Amphotericin B (Ampho B) decreases the stiffness of IFITM3-containing membranes and prevents virus inhibition. These data provide significant new insight into the mechanism by which IFITM3 inhibits the fusion of diverse pathogenic viruses and reveal that oligomerization is a shared requirement for the distinct anti-fusion functions performed by homologs IFITM3 and PRRT2.

## Results

### Evolution-guided identification of a putative oligomerization motif within CD225 domains

A phylogenetic tree of CD225 domains found in the *IFITM* family (consisting of *IFITM1, IFITM2, IFITM3, IFITM5*, and *IFITM10* in humans) and other CD225 proteins (human *PRRT2* and *TUSC5*) are indicative of common ancestry **(Figure 1A)**. While it has been suggested that IFITM3 may adopt multiple topologies [17, 48], experimental evidence indicates that IFITM3 is a type II transmembrane protein characterized by the presence of a cytoplasmic-facing amino terminus, the CD225 domain consisting of a hydrophobic intramembrane (IM) domain, a cytoplasmic intracellular loop (CIL), and a hydrophobic transmembrane (TM) domain, and a very short carboxy terminus facing the vesicle lumen or extracellular space [49, 50]. PRRT2 is thought to adopt a similar topology in membranes [51]. We used Protter, an interactive application that maps annotated and predicted protein sequence features onto the transmembrane topology of proteins [52], to visualize IFITM3 and PRRT2. The two proteins exhibit a similar predicted topology consisting of dual hydrophobic domains and a CIL, but the amino terminus is considerably longer in PRRT2 **(Figure 1B and Supplemental Figure 1A)**. When comparing topologies and the protein alignment **(Figure 1C)**, we noticed the presence of a GxxxG motif in the CIL of IFITM3 and PRRT2 **(Figure 1B and 1C)** which is intact in several other IFITM and CD225 proteins [5]. The GxxxG motif, also known as a glycine zipper, is frequently associated with dimerization of membrane proteins [53, 54]. Most often shown to mediate pairing of hydrophobic transmembrane helices within a bilayer, the motif has also been described to drive oligomerization from cytoplasmic loops or linkers [55]. The GxxxG motif is conserved in *IFITM3* of vertebrates, indicating that it may play an important functional role **(Figure 1D)**.

**Figure 1:**
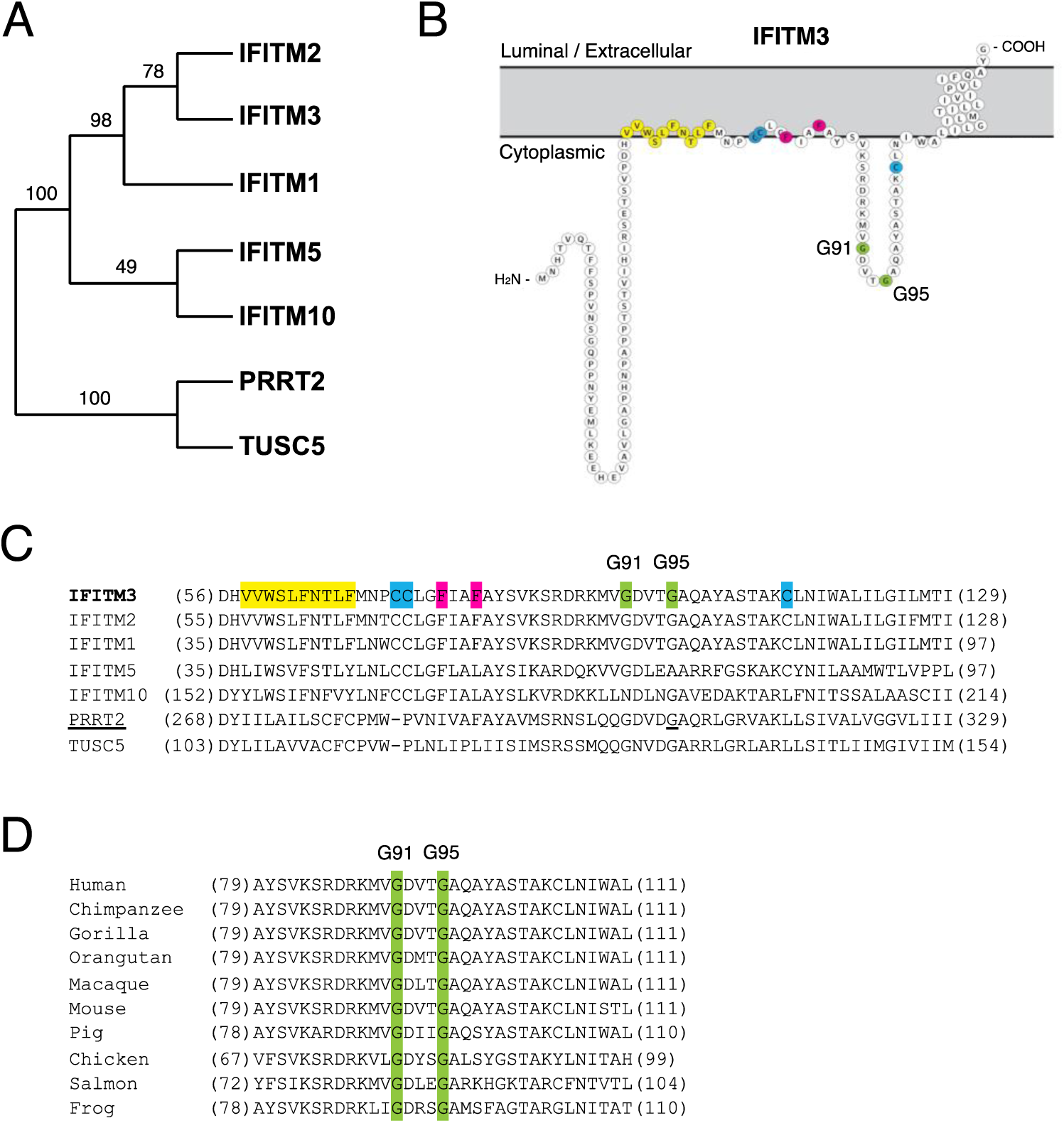
Identification of a GxxxG motif in CD225 domains. (A) A phylogenetic tree of CD225 amino acid sequences from IFITM and other CD225 proteins was reconstructed by maximum likelihood and presented as a cladogram. Reliability of internal branch topology was assessed by bootstrapping (100 replicates). (B) Schematic representation of the membrane topology of IFITM3 made with Protter. Residues corresponding to the amphipathic helix (yellow), palmitoylated cysteines (blue), phenylalanines purported to regulate oligomerization (red), and the glycines of the GxxxG motif (green) are indicated. (C) A partial amino acid alignment of CD225 domains from IFITM proteins, PRRT2, and TUSC5. Color codes are included as in (B). The position of the polymorphic glycine in PRRT2 associated with neurological disease (G305W) is underlined. (D) A partial amino acid alignment of IFITM3 orthologs in vertebrates. Conserved glycines in the GxxxG motif are indicated in green.

### Glycine-95 is important for the antiviral functions of IFITM3

We generated FLAG-tagged IFITM3 mutants in which glycine-91 and glycine-95 were changed to leucine (G91L and G95L), following an example set by characterization of the GxxxG motif in the human folate transporter [56]. We also produced a G95W mutant because a rare single nucleotide polymorphism known as rs779445843 gives rise to missense mutations (G95W/R) in human populations [57, 58]. Furthermore, G95W in IFITM3 is analogous to a disease-associated polymorphism in PRRT2 (G305W) which results in loss of function [44]. Upon transfection into human cells, IFITM3 containing G95L or G95W exhibited steady-state protein levels similar to wild-type (WT), but this was not the case for G91A **(Supplemental Figure 2A)**. This finding may explain why a previous study failed to express an IFITM3 mutant containing a stretch of alanines between positions 90 and 95 [13, 34]. Therefore, we focused on G95L and G95W for further functional characterization. The subcellular localization of IFITM3 WT and G95L was found to be similar when confocal immunofluorescence microscopy was performed. As shown previously [3, 59, 60], protein was detected in early endosomes, late endosomes, and at the plasma membrane **(Supplemental Figure 2B)**. We generated HEK293T cell lines stably expressing FLAG-tagged IFITM3 WT and mutants to assess antiviral function **(Figure 2A)**. Challenge of cell lines with IAV demonstrated that, while IFITM3 WT strongly protected cells from infection, the G95L or G95W mutations resulted in a considerable loss of virus restriction **(Figure 2B)**. When we tested antiviral function following transient transfection into HEK293T, we found that G95L or G95W resulted in a complete loss of IAV restriction **(Figure 2C and Supplemental Figure 2A)**. To confirm that IFITM3 targets the entry step of IAV, we challenged cells with HIV-1 pseudotyped with hemagglutinin (HA, a type 1 viral fusogen) and found that G95L results in a severe loss of pseudovirus restriction **(Figure 2D)**. The G95L and G95W mutations also abrogated restriction of HIV-1 pseudotyped with VSV glycoprotein (a type 3 viral fusogen) in an assay for virus-cell fusion **(Figure 2E)**. Therefore, a rational approach identified glycine-95 to be essential for broad inhibition of virus entry by IFITM3.

**Figure 2:**
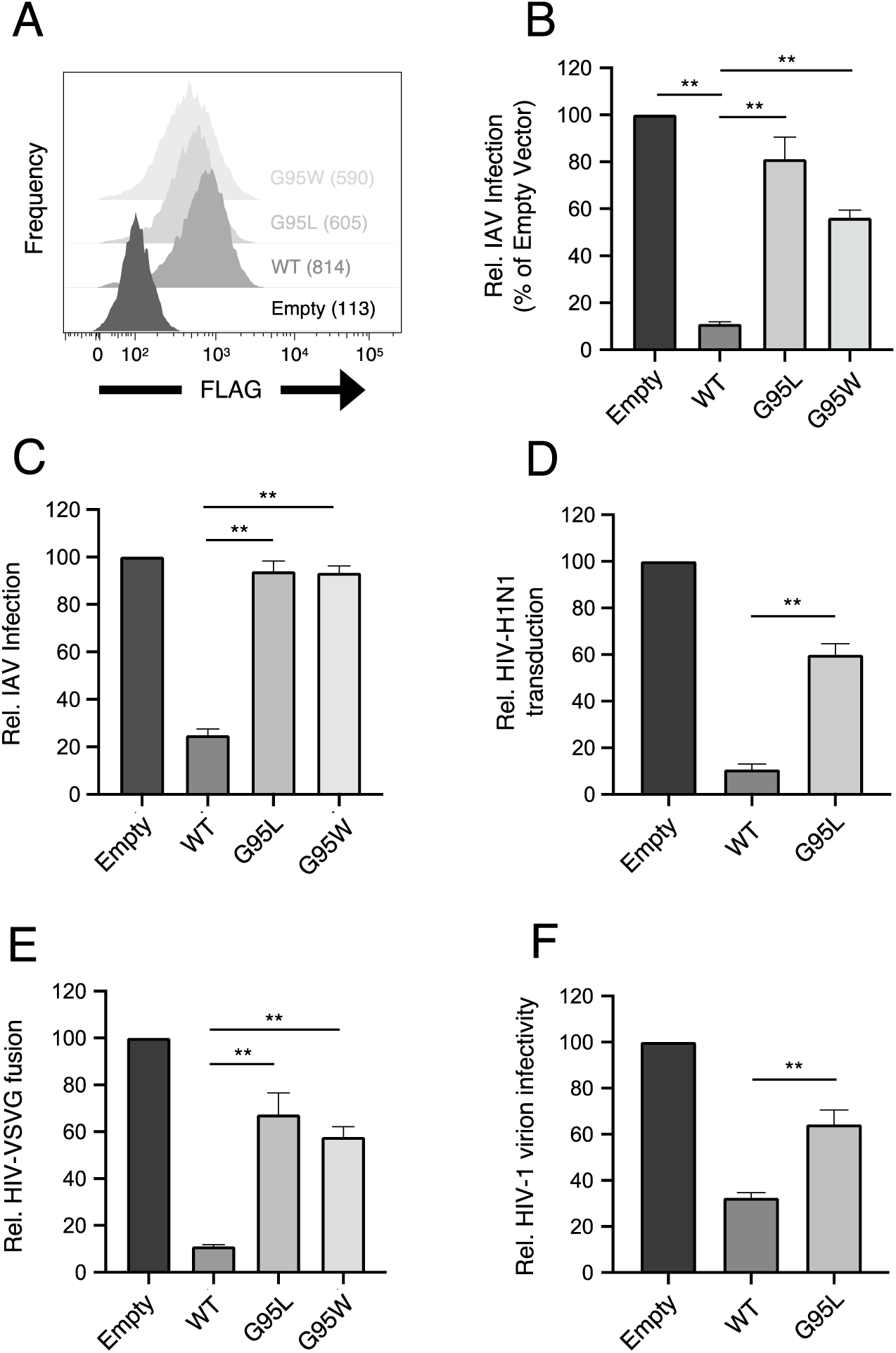
The role of glycine-95 in the antiviral functions of IFITM3. (A) HEK293T cells stably transfected with empty pQCXIP, WT IFITM3-FLAG, or the indicated mutants were fixed, stained with anti-FLAG antibody and assessed by flow cytometry. FLAG levels are displayed as histograms and values corresponding to mean fluorescence intensity are indicated. (B) Cell lines from (A) were challenged with IAV PR8 strain (MOI of 0.1) and fixed, stained with an anti-nucleoprotein antibody, and assessed by flow cytometry at 18 hours post-infection. (C) HEK293T were transiently transfected with 1.5 micrograms of empty pQCXIP, WT IFITM3-FLAG, or the indicated mutants and, 48 hours later, challenged with IAV PR8 strain. (D) Stably transfected HEK293T cells were challenged with replication-incompetent HIV-1 pseudotyped with hemagglutinin and neuraminidase from H1N1 IAV. Infection was scored by measurement of GFP+ cells by flow cytometry at 48 hours post-infection. (E) Stably transfected HEK293T were challenged with replication-incompetent HIV-1 pseudotyped with glycoprotein from VSV. Virus-cell fusion was assessed using the BlaM-Vpr assay at 2.5 hours post-virus addition. (F) HEK293T were co-transfected with HIV-1 molecular clone NL4.3 and empty pQCXIP, WT IFITM3-FLAG, or the indicated mutant. Virus-containing supernatants were harvested at 48 hours post-transfection and subjected to ultracentrifugation over sucrose pellets. Virion content was quantified by p24 CA ELISA and 50 ng p24 equivalent was added to TZM.bl cells for infectivity measurements. TZM.bl were fixed at 48 hours post-infection, stained with anti-CA antibody, and infection was assessed by flow cytometry. Results represent the mean of 3-8 independent experiments and are normalized to empty vector (set to 100%). Error bars indicate standard error. Rel., relative. Statistical analysis was performed using one-way ANOVA. *, P < 0.05; **, P < 0.001.

We and others have previously demonstrated that, in addition to preventing virus entry into naïve target cells, IFITM3 performs another antiviral function in virus-producing (infected) cells by incorporating into virions, reducing viral glycoprotein abundance and function, and reducing the fusogenic potential of virions [25-29, 32]. Therefore, we also tested the relative impact of IFITM3 WT and IFITM3 G95L on HIV-1 virion infectivity. IFITM3 WT reduced the infectivity of HIV-1 and was accompanied by a defect in viral Envelope (Env) incorporation into virions: processing of the Env gp160 precursor (as determined by gp120/gp160 ratio) was decreased, resulting in relatively low levels of gp120 and gp41, consistent with previous work [26, 28, 61]. By comparison, IFITM3 G95L exhibited reduced activity against HIV-1 infectivity and impacted Env processing/incorporation to a lesser extent **(Figure 2F and Supplemental Figure 2C)**. Furthermore, IFITM3 G95L itself was incorporated into HIV-1 virions about 50% less than WT IFITM3 **(Supplemental Figure 2C)**. Therefore, the dual antiviral functions performed by IFITM3 (early-stage inhibition of virus entry and late-stage inhibition of virion infectivity) are regulated by glycine-95.

### ^91^GxxxG^95^ regulates oligomerization of IFITM3 in living cells and bulk lysates

Based on its position within a conserved GxxxG motif, we used FRET to assess the role played by glycine-95 in the oligomerization of IFITM3. We constructed IFITM3 fused with yellow fluorescent protein (YFP) or mCherry to create FRET pairs and to perform FLIM. The measurement of fluorescence lifetimes allows for measurements that are independent of fluorophore expression level or diffusion rate [62]. In this framework, excitation of YFP (donor) results in energy transfer to mCherry (acceptor) when a molecular interaction brings the pair into close proximity. Co-transfection of mCherry and IFITM3-YFP served to establish background FRET values. By comparison, WT IFITM3-YFP and WT IFITM3-mCherry co-transfection resulted in high levels of detectable FRET **(Figure 3A and 3B)**. However, introduction of the G95L mutation significantly reduced FRET, suggesting that glycine-95 is important for oligomerization. FRET was also measured for heterologous pairs (combining WT IFITM3-YFP and G95L-mCherry, and vice versa) and the results were indicative of an intermediate degree of oligomerization **(Figure 3B)**. To complement our FRET analysis, we also analyzed fluorescence lifetimes of the donor (YFP). Co-transfection of WT IFITM3-YFP and WT IFITM3-mCherry resulted in significant decreases in YFP lifetimes, consistent with IFITM3 oligomerization **(Figure 3A and 3C)**. However, we did not observe decreases in YFP lifetimes when G95L-YFP and G95L-mCherry were examined. To ensure that conclusions drawn from this imaging approach are relevant to IFITM3-mediated antiviral function, we tested the ability for fluorescently-tagged IFITM3 to inhibit IAV. WT IFITM3-mCherry exhibited nearly the same antiviral potency as WT IFITM3-FLAG while WT IFITM3-YFP was slightly less potent. Moreover, versions of fluorescently-tagged IFITM3 encoding G95L exhibited significantly less antiviral activity than their WT counterparts **(Supplemental Figure 3A)**. Using the same FRET-based approach, we found that the G91L mutation partially resulted in higher YFP lifetimes relative to WT IFITM3, indicating that G91L also impairs oligomerization of IFITM3 **(Supplemental Figure 3B)**. Overall, these data provide the first indication that glycine-91 and glycine-95 are important for IFITM3 oligomerization. To quantitatively resolve the specific oligomeric state of IFITM3 under these conditions, we performed a Number and Brightness analysis [63, 64]. This approach is a fluorescence microscopy method capable of measuring the apparent average number of molecules and their brightness in each pixel over time, with brightness being proportional to oligomeric state. Analysis was restricted to IFITM3-mCherry during its diffusion in and out of a static membranous compartment (the plasma membrane). On average, WT IFITM3-mCherry was found to be 2.14 times brighter than mCherry monomers [65] **(Figure 3D)**. In contrast, G95L-mCherry brightness was not significantly different than mCherry monomers (averaging 1.30 times brighter). Furthermore, relative to mCherry monomers, both WT IFITM3-mCherry and G95L-mCherry formed assemblies that were up to five times brighter, but WT IFITM3-mCherry demonstrated a greater propensity to form these higher-order oligomers **(Figure 3D)**. These data suggest that fluorescently-tagged IFITM3 forms dimers and higher-order oligomers in membranes in a glycine-91- and glycine-95-dependent fashion.

**Figure 3:**
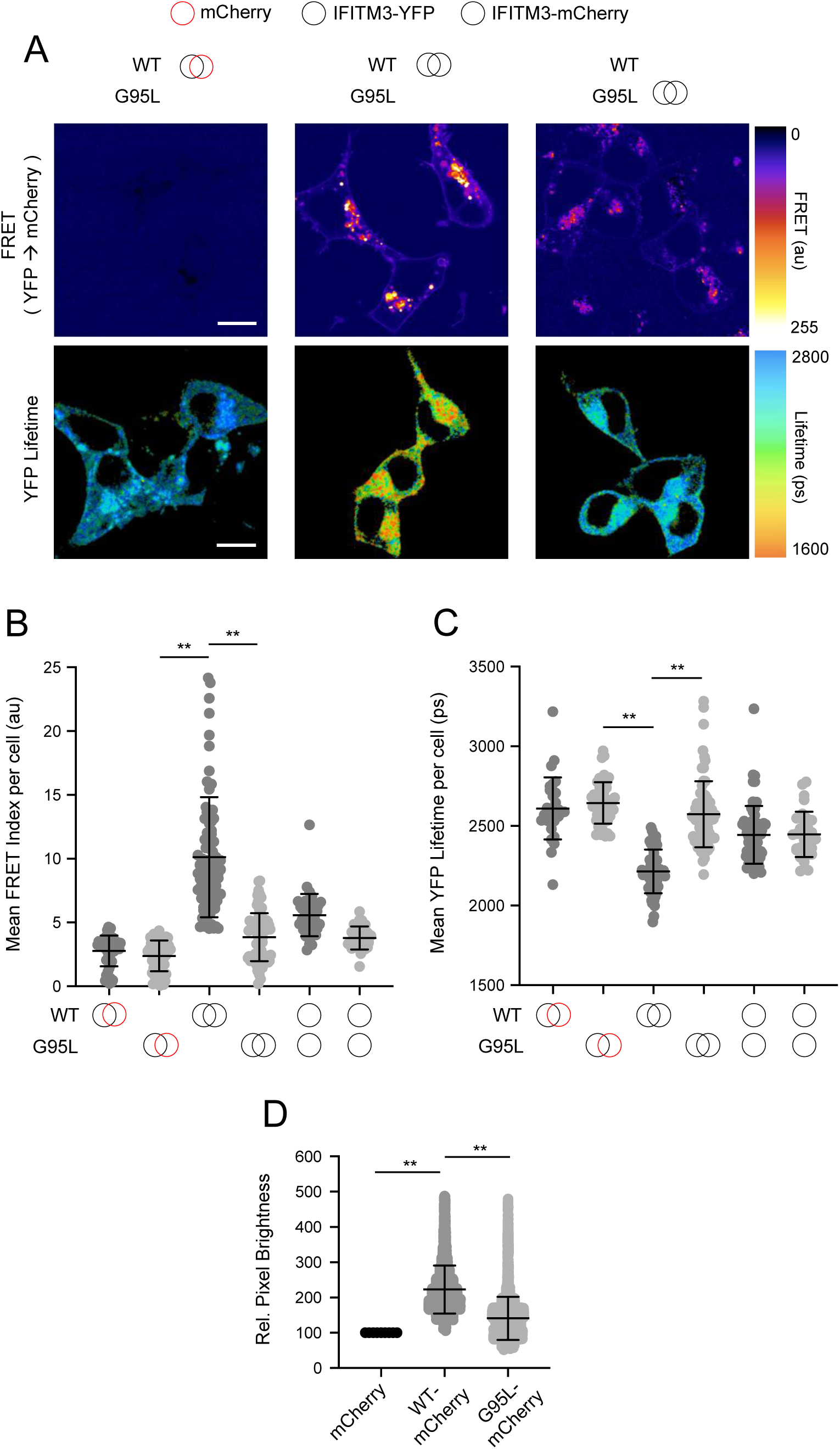
^91^GxxxG^95^ regulates oligomerization of IFITM3 in living cells. (A) HEK293T were transiently co-transfected with IFITM3-YFP and mCherry or IFITM3-YFP and IFITM3-mCherry. Constructs encoded WT IFITM3 or IFITM3 G95L. FRET-FLIM measurements were made, and images of FRET signal and YFP lifetimes are representative of 12-20 captured images per condition. (B) Whole-cell FRET analysis was performed on a minimum of 50 cells per condition and the results of three independent experiments were pooled. Dots correspond to individual cells. (C) Whole-cell YFP lifetimes were measured on a minimum of 50 cells per condition and the results of three independent experiments were pooled. Dots correspond to individual cells. (D) Number and Brightness analysis was performed on monomeric mCherry and IFITM3-mCherry (either WT or G95L). Empty red circles are used to depict mCherry, filled red circles are used to depict IFITM3-mCherry (either WT or G95L), and filled yellow circles are used to depict IFITM3-YFP (either WT or G95L). Error bars indicate standard deviation. Statistical analysis was performed using one-way ANOVA. *, P < 0.05; **, P < 0.001. Scale bars, 10 μm. Ps, picoseconds; Au, arbitrary units; Rel., relative.

In parallel to our studies of IFITM3 oligomerization in single, living cells, we assayed the ability of IFITM3 pairs tagged with FLAG or myc to co-immunoprecipitate from bulk cell lysates. HEK293T were co-transfected with IFITM3-FLAG and IFITM3-myc followed by FLAG immunoprecipitation, SDS-PAGE, and quantitative immunoblotting. We found that WT IFITM3-myc readily pulled down with WT IFITM3-FLAG, while pull down of G95L-myc with G95L-FLAG was diminished by approximately 50% **(Figure 4A and 4B)**. Therefore, membrane-extracted IFITM3 forms oligomers, but the G95L mutation reduces oligomerization. We then performed blue native PAGE and immunoblotting to assess the oligomeric state of IFITM3 under non-denaturing conditions. Two populations of IFITM3 oligomers, exhibiting sizes of approximately 300 and 480 kilodaltons, were readily observed for WT and, to a lesser extent, IFITM3 G95L. The largest (480 kilodaltons) was nearly absent for IFITM3 G95L **(Figure 4C)**. In contrast to the Number and Brightness analysis **(Figure 3D)**, we did not observe dimers, which may result from the conditions under which blue native PAGE was performed here **(Supplemental Figure 4A)**. Nonetheless, the fact that IFITM3 G95L exhibited a reduced potential for higher-order oligomer formation using this technique is consistent with our experiments using living cells or denaturing conditions. Therefore, glycine-95 is necessary for efficient IFITM3 oligomer formation and oligomerization is necessary for its antiviral function.

**Figure 4:**
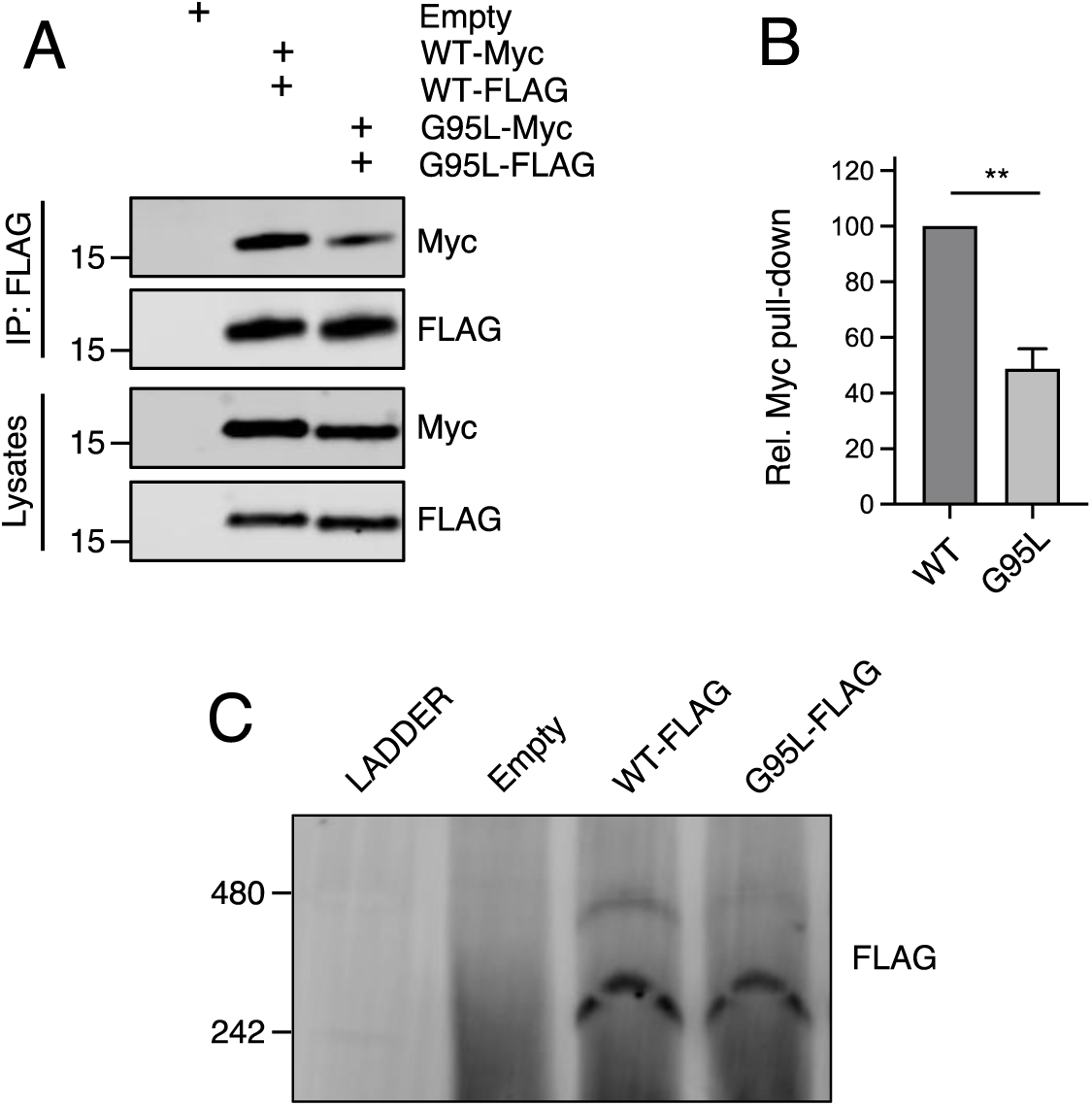
glycine-95 regulates oligomerization of IFITM3 in bulk cell lysates. (A) HEK293T were transiently transfected with empty pQCXIP or the following pairs: IFITM3-FLAG and IFITM3-myc or G95L-FLAG and G95L-myc. Whole cell lysates were produced under mildly denaturing conditions and immunoprecipitation (IP) using anti-FLAG antibody was performed. IP fractions and volumes of whole cell lysates were subjected to SDS-PAGE and Western blot analysis. Immunoblotting was performed with anti-FLAG and anti-myc. Number and tick marks indicate size (kilodaltons) and position of protein standards in ladder. (B) Levels of IFITM3-myc (either WT or G95L) co-immunoprecipitated by anti-FLAG IP were quantified from conditions in (A). Results represent the mean of three independent experiments. Error bars indicate standard error. (C) HEK293T were transiently transfected with empty pQCXIP, WT IFITM3-FLAG or G95L-FLAG. Cell lysates were produced with 1% digitonin and blue native PAGE was performed, followed by immunoblotting with anti-FLAG. Number and tick marks indicate size (kilodaltons) and position of protein standards in ladder. Statistical analysis was performed using student’s T test. *, P < 0.05; **, P < 0.001.

### Membrane order is increased by IFITM3 oligomers and is a correlate of antiviral function

While the precise mechanism by which IFITM3 inhibits virus-cell fusion remains unresolved, a salient phenotype of IFITM protein expression is increased membrane order (reduced membrane fluidity) [17, 34]. Therefore, we leveraged our loss-of-function mutant to directly test whether membrane order is functionally associated with inhibition of virus entry by IFITM3. Previous reports of membrane order enhancement by IFITM family members involved the use of a cell-permeable dye known as Laurdan [66]. Here, we assessed membrane order using a recently described sensor known as fluorescent lipid tension reporter (FliptR) [67-69]. FliptR is a planarizable push-pull probe that incorporates efficiently into artificial and living cell membranes and whose fluorescence parameters change upon alterations in local lipid packing (order). Specifically, FliptR responds to increasing membrane order at the plasma membrane and at endomembranes by planarization, leading to longer fluorescence lifetimes detected by FLIM [67, 70]. Using HEK293T stably expressing FLAG-IFITM3, we found that WT IFITM3 expression led to significantly increased membrane order, consistent with previous reports [14, 17] **(Figure 5A)**. In fact, the IFITM3-induced enhancement of membrane order was similar to that achieved by addition of soluble cholesterol, while cholesterol depletion with methyl-beta-cyclodextrin resulted in profound decreases in membrane order **(Supplemental Figure 5A)**. In contrast to WT, cells expressing IFITM3 G95L did not exhibit increased membrane order **(Figure 5A)**. These data indicate that membrane order enhancement tracks with a functionally-competent form of IFITM3 but not with a loss-of-function mutant. To further probe the functional importance of membrane order in the antiviral mechanism of IFITM3, we performed experiments in the presence of Ampho B. This antimycotic polyene compound was previously reported to overcome the antiviral activity of IFITM3, rendering cells stably expressing IFITM3 fully permissive to IAV [14]. However, it was unknown how Ampho B counteracts the effects of IFITM3. When we added Ampho B to cells expressing WT IFITM3, we no longer observed increased membrane order **(Figure 5A)**. Furthermore, in identically-treated cells, Ampho B prevented restriction of HA-mediated virus entry by IFITM3 **(Figure 5B)**. These findings show, for the first time, that the capacity for Ampho B to overcome the antiviral activity of IFITM3 is linked to its ability to decrease membrane order. Therefore, the use of Ampho B as an interrogative tool reinforced the role played by membrane order in the antiviral mechanism of IFITM3. Overall, these data strongly suggest that IFITM3-mediated antiviral activity occurs through oligomerization-dependent membrane stiffening.

**Figure 5:**
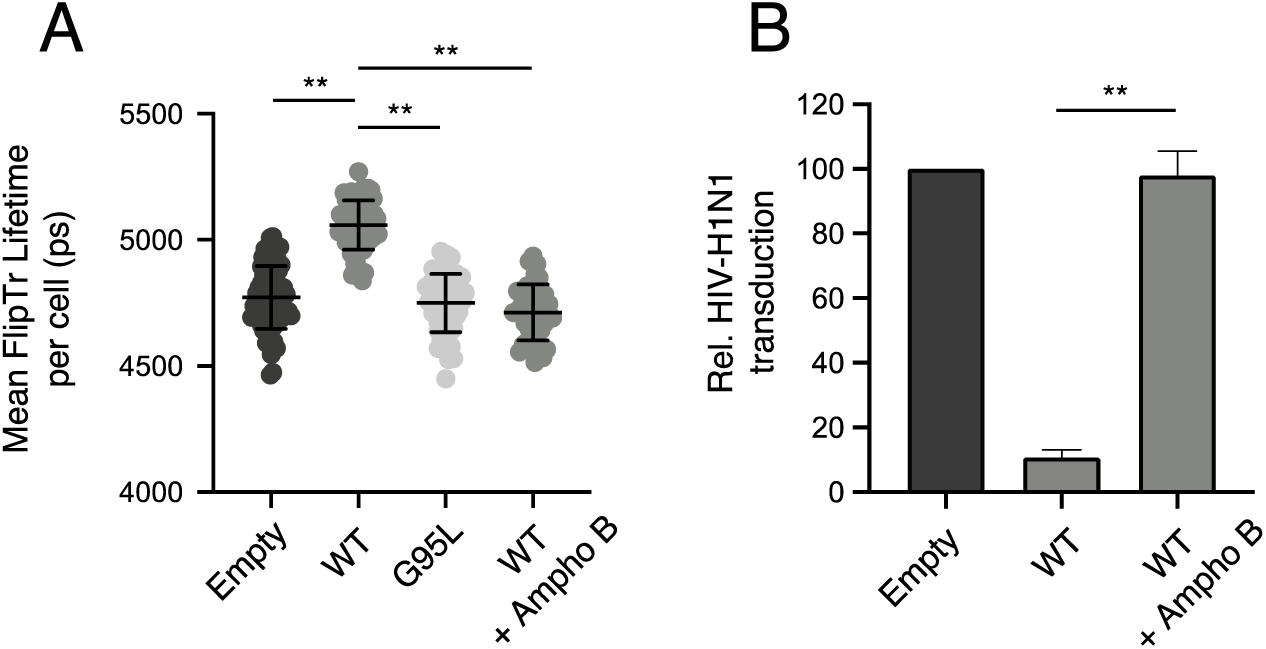
Membrane order is increased by IFITM3 oligomers in an Ampho B-sensitive manner. (A) HEK293T cells stably transfected with empty pQCXIP, WT IFITM3-FLAG, or G95L-FLAG were stained with 1 μM FliptR for 5 minutes and imaged by FLIM. In the condition indicated, 1 μM amphotericin B was added to cells for one hour and washed away prior to addition of FliptR and imaging. The whole-cell mean fluorescence lifetime (τ), in addition to individual component lifetimes (long and short lifetimes, τ_1_ and τ_2_), were calculated using Symphotime for a minimum of 40 cells per condition and τ_1_ from three independent experiments were pooled and plotted. Dots correspond to individual cells. Error bars indicate standard deviation. (B) HEK293T cells stably transfected with empty pQCXIP, WT IFITM3-FLAG, or G95L-FLAG were challenged with HIV pseudotyped with hemagglutinin and neuraminidase proteins from IAV WSN strain at a MOI of 1. In the condition indicated, 1 μM amphotericin B was added to cells for one hour prior to virus addition. Cells were fixed at 48 hours post-infection and infection was scored by GFP expression using flow cytometry. Statistical analysis was performed using one-way ANOVA. *, P < 0.05; **, P < 0.001.

### Disease-associated G305W impairs oligomerization of PRRT2

Since the GxxxG motif is a shared feature of IFITM3 and PRRT2 and that naturally-occurring variation at glycine-305 of PRRT2 is predictive of neurological disease, we assessed the oligomerization capacity of WT and mutant PRRT2. As in Figure 3, we constructed PRRT2 fused with YFP or mCherry to create FRET pairs. We observed that co-transfection of PRRT2-YFP and PRRT2-mCherry resulted in significant FRET, demonstrating that PRRT2 oligomerizes in living cell membranes **(Figure 6A)**. However, FRET was significantly reduced for pairs containing G305W (G305W-YFP and G305W-mCherry) **(Figure 6A)**. Furthermore, pairs containing G305W did not exhibit loss in YFP lifetimes relative to their WT counterparts **(Figure 6B)**. These data suggest that mutation of glycine-305 in PRRT2 results in loss of protein oligomerization. Therefore, the divergent functions played by homologues IFITM3 and PRRT2 in the regulation of fusion processes are controlled by a common determinant.

**Figure 6:**
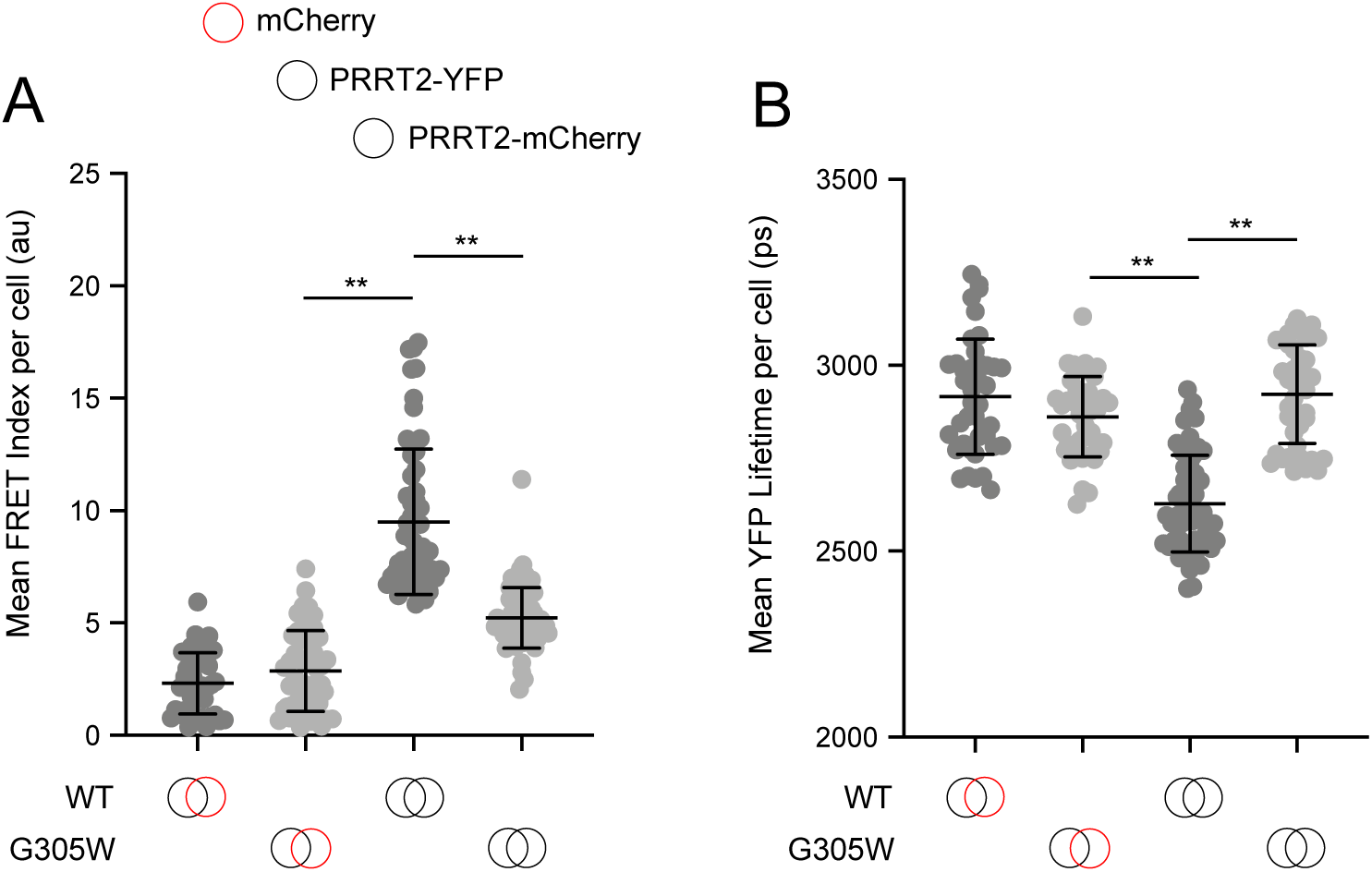
Disease-associated G305W disrupts the oligomerization of PRRT2 in living cells. (A) HEK293T were transiently co-transfected with PRRT2-YFP and mCherry or PRRT2-YFP and PRRT2-mCherry. Constructs encoded WT PRRT2 or PRRT2 G305W. Whole-cell FRET analysis was performed on a minimum of 50 cells per condition and the results of three independent experiments were pooled. Dots correspond to individual cells. (B) Whole-cell YFP lifetimes were measured on a minimum of 50 cells per condition and the results of three independent experiments were pooled. Dots correspond to individual cells. Empty red circles are used to depict mCherry, filled red circles are used to depict PRRT2-mCherry (either WT or G305W), and filled yellow circles are used to depict PRRT2-YFP (either WT or G305W). Error bars indicate standard deviation. Statistical analysis was performed using one-way ANOVA. *, P < 0.05; **, P < 0.001.

## Discussion

The physiological importance of IFITM3 in the control of many virus infections in vivo is becoming increasingly apparent [71, 72]. While it has been proposed that membrane remodeling by IFITM3, at the level of membrane order and curvature, protects host cells from virus invasion, functional proof was lacking. However, two recent developments provided a glimpse of how (and when) IFITM3 inhibits virus-cell membrane fusion. First, the identification of an amphipathic helix located in the IM domain of IFITM3 provided a rational explanation for how membrane stiffening and/or bending may occur [22]. Second, IFITM3 has been observed to intercept vesicles carrying inbound virions and to restrict their release into the cytoplasm, while viruses that are insensitive to IFITM3 evade its encounter [31, 73]. Together with the previous demonstrations that the subcellular localization of IFITM3 determines its specificity and potency [30, 74, 75], a “proximity-based” mechanism presents itself in which IFITM3 interacts with and modifies host membranes needed by some viruses to fuse with cells. Importantly, this model accommodates the antiviral effect of IFITM3 inside viral lipid bilayers as well [25, 26, 28, 31, 32].

Here, we provide evidence that an additional determinant (protein oligomerization) plays a crucial role within this mechanistic framework. Previously, an alanine scanning approach led to the identification of two residues (phenylalanine-75 and phenylalanine-78) that were important for IFITM3 oligomerization in cell lysates [34]. Importantly, this study did not functionally test a role for glycine-91 or glycine-95 in the oligomerization of IFITM3 because alanine mutagenesis overlapping these residues led to loss of stable protein expression [13, 34]. While it has been confirmed that F75A and F78A mutations disrupt antiviral activity [31, 76], IFITM3 oligomerization was not impacted by these mutations when assessed by FRET in living cells [35]. It remains possible that these two phenylalanines cooperate with the GxxxG motif to mediate IFITM3 oligomerization. However, our data shows that G95L reduces the antiviral potential of IFITM3 as well as its capacity to increase membrane order (reduce membrane fluidity). Furthermore, we show that a compound previously found to abrogate the antiviral function of IFITM3, Ampho B [14, 31], decreases membrane order in IFITM3-expressing cells. Even though the exact mechanism by which Ampho B impacts mammalian membranes is unclear [14, 77], these results identify that the membrane stiffening property of IFITM3 is a strict correlate of its antiviral functions in cells and, most likely, in virions. We found that the oligomerization-defective G95L mutant lessened the impact of virion-associated IFITM3 on HIV-1, suggesting that IFITM3 oligomers are needed to maximally reduce virion infectivity.

It will be important to assess how oligomerization-defective mutants of IFITM3 (G95L) impact membrane curvature, another reported consequence of ectopic IFITM3 expression in cells [17]. It has been shown that protein oligomerization of the transmembrane protein Mic10 is essential for its capacity to induce curvature in mitochondrial membranes [78]. Furthermore, changes in membrane order are often accompanied by changes in membrane curvature [18-20]. It is possible that these two alterations to host membranes underlie the restriction of virus fusion by IFITM3. Moreover, amphipathic helices are characterized to interact with and bend membranes [24]. However, the impact that the amphipathic helix of IFITM3 has on membrane order and curvature has not been assessed. Regardless of how the amphipathic helix is responsible for inhibition of virus fusion, our data raise the possibility that oligomerization “activates” the antiviral potential of the amphipathic helix. It is possible that local insertion of multiple amphipathic helices into stretches of membrane is required for inhibition of virus fusion, and IFITM3 oligomers provide a means to fulfill that requirement.

In addition to mediating dimerization or oligomerization of transmembrane proteins (homomultimerization), GxxxG motifs have also been described to affect the propensity for interaction with other proteins (heteromultimerization) [53, 79]. Therefore, glycine-95 may also govern which proteins IFITM3 interacts with and to what extent. IFITM3 has been described to bind with IFITM1 and IFITM2, and it is interesting to consider how IFITM heteromultimers may contribute to antiviral protection of the cell [34]. Furthermore, other host proteins have been described to interact, directly or indirectly, with IFITM3. This list includes cholesterol trafficking regulator VAPA and the metalloproteinase ZMPSTE24, two proteins that have been described as essential cofactors for the antiviral effects of IFITM3 [10, 80]. Since the former is associated with the tendency for IFITM3 to cause cholesterol overload in endosomes, the G95L loss-of-function mutant could be used to rule in or rule out VAPA and cholesterol as players in the antiviral mechanism of IFITM3.

While glycine-95 is critical for the oligomerization and anti-fusion activity of IFITM3, we show that the homologous site in PRRT2, glycine-305, regulates its oligomerization as well. The naturally-occurring G305W/R mutations found in patients with neurological dysfunction are known to disrupt PRRT2 activity, and our results here provide novel insight into how loss-of-function occurs. Therefore, the presence of a shared GxxxG motif in IFITM3, PRRT2, and some other CD225 family members suggests that an ancestral CD225-containing protein performed an unknown function that required oligomerization. We wonder whether all CD225 proteins play roles in regulating membrane fusion processes in cells—only time and further experiments will tell. However, just as glycine-95 in IFITM3 may mediate both homo-and heteromultimerization, it has been reported that PRRT2 interacts with cellular fusogens known as soluble N-ethylmaleimide-sensitive factor attachment protein receptors (SNAREs) in a glycine-305-dependent manner [44]. Our findings raise the possibility that oligomerization of CD225 proteins results in alteration of protein architecture and the display of novel docking sites for protein-protein interactions, allowing for expansion of their functional repertoire.

## Materials & Methods

### Sequence alignments and phylogenetic analysis

Protein sequences for CD225 proteins (including IFITM proteins, PRRT2, and TUSC5) were retrieved from UniProt and multi-sequence alignments were performed with MUSCLE. Gaps were removed with Gblocks and phylogenetic reconstruction by maximum likelihood was performed with PhyML, using the WAG substitution model. Branch confidence was assessed using the bootstrapping method (100 replicates). Graphic representation of the tree as a cladogram was performed with TreeDyn.

### Cell lines and plasmids

HEK293T (ATCC: CRL-3216) and TZM-bl (NIH AIDS Reagent Resource: 8129) and any derivatives produced in this study were cultivated at 37°C and 5% C02 in DMEM (Gibco) complemented with 10% fetal bovine serum (Hyclone) and 1% penicillin-streptomycin (Gibco) and regularly passaged with the aid of Trypsin-EDTA 0.05% (Gibco). Retroviral pQCXIP vectors encoding IFITM3 fused with amino-terminal FLAG were previously described [30]. Retroviral pQCXIP vectors encoding IFITM3 fused with amino-terminal myc were produced by appending myc to the open reading frame of IFITM3 with PCR and cloning into BamH1/EcoR1 sites. WT PRRT2 fused with carboxy-terminal YFP or mCherry was produced by Integrated DNA Technologies. The G91L, G95L, and G95W mutations in IFITM3 and the G305W mutations in PRRT2 were introduced by site-directed mutagenesis (QuikChange Lightning) or by ligation-independent cloning. HEK293T cell lines stably expressing pQCXIP plasmids were produced by transfecting 250,000 cells in a 12-well plate with 0.8 μg DNA using Lipofectamine2000 (Invitrogen) and performing puromycin selection at a concentration of 10 μg/mL for two weeks.

### Virus productions, infections and virus-cell fusion assay

Influenza A Virus [A/PR/8/34 (PR8), H1N1] supplied as clarified allantoic fluid was purchased from Charles River Laboratories. Infectious virus titers were calculated using a flow cytometry-based method in HEK293T cells [81], and infections were performed as follows: HEK293T cells (either stably expressing or transiently transfected with 1.5 μg empty pQCXIP or WT IFITM3 or the indicated mutant) were seeded in 24-well plates (50,000 per well) overnight and overlaid with indicated amounts of virus diluted in 225 μL of complemented DMEM for approximately 18 hours. Cells were washed with 1X PBS, detached with Trypsin-EDTA, fixed/permeabilized with Cytofix/Cytoperm (BD), immunostained with anti-IAV NP (AA5H; Abcam), and analyzed on a LSRFortessa flow cytometer (BD). Replication-incompetent HIV-1 pseudotyped with IAV WSN HA and NA was produced by transfecting HEK293T with 2 μg pR8ΔEnv, 1 μg pcRev (NIH AIDS Reagent Resource: 11415), 3 μg Gag-GFP, 1.5 μg of hemagglutinin, and 1.5 μg of neuraminidase from IAV WSN strain, H1N1 (gifts from G. Melikyan). Replication-competent HIV-1 was produced by transfecting HEK293T with 15 μg pNL4-3 and 5 μg empty pQCXIP or pQCXIP encoding IFITM3 WT or the indicated mutant fused with amino-terminal FLAG. Replication-incompetent HIV-1 pseudotyped with VSV-G for virus-cell fusion assays was produced by transfecting HEK293T with 15 μg pNL4-3ΔEnv, 5 μg pCMV4-BlaM-Vpr, and 5 μg pMD2.G (VSV-G). Transfections were performed using the calcium-phosphate method. Briefly, six million HEK293T were seeded in a T75 flask. Plasmid DNA was mixed with sterile H_2_0, CaCl_2_, and Tris-EDTA (TE) buffer, and the totality was combined with Hepes-buffered saline (HBS). The transfection volume was added dropwise, and cells were incubated at 37°C for 48 hours. Supernatants were clarified by centrifugation, passed through a 0.45-μm filter, and concentrated by ultracentrifugation through a 20% sucrose cushion at 25000 x g for 1 hour at 4°C. Lentivirus titers were measured using an HIV-1 p24 ELISA kit (XpressBio). To measure infectivity of virus supernatants, 50 ng p24 equivalent volumes were added to TZM-bl cells and cells were fixed/permeabilized with Cytofix/Cytoperm (BD) at 48 hours post-infection, immunostained with anti-Gag KC57-FITC (BD), and analyzed by flow cytometry. To measure virus-cell fusion, 100-300 ng p24 equivalent HIV-1-VSV-G produced with pCMV4-BlaM-Vpr was added to HEK293T cells stably expressing WT IFITM3 or the indicated mutant for 2.5 hours at 37°C. Cells were washed with CO_2_-independent medium and incubated with CCF2/AM mix containing probenecid for one hour at room temperature in the dark. Cells were washed with cold 1X PBS, fixed/permeabilized with Cytofix/Cytoperm (BD), and analyzed on a LSRII flow cytometer.

### Confocal immunofluorescence microscopy

HEK293T cells were seeded in μ-slide 8 well chambers (Ibidi) (30,000 per well) overnight and transfected with 0.2 μg EEA1-GFP (Addgene: 42307) and 0.2 μg empty pQCXIP or pQCXIP-WT IFITM3 or the indicated mutant fused with amino-terminal FLAG using Lipofectamine2000 (Invitrogen). At 48 hours post-transfection, cells were fixed/permeabilized with Cytofix/Cytoperm and immunostained with anti-CD63 antibody (MX-49, sc-5275; Santa Cruz Biotechnology) and anti-IFITM3 antibody (EPR5242; Abcam). Cells were imaged using the Leica TCS SP8 confocal microscope with a 63X objective and oil immersion and analysis was performed in Fiji (ImageJ).

### Immunoprecipitation, SDS-PAGE, and Western blot analysis

HEK293T cells were transfected with 1.5 μg empty pQCXIP or a combination of 0.75 μg pQCXIP-FLAG-IFITM3 and 0.75 μg pQCXIP-myc-IFITM3 (encoding either WT or G95L). At 48 hours post-transfection, cells were lysed in a buffer containing 0.5% IPEGAL (Sigma), 50 mM Tris (pH 7.4), 150 mM NaCl, and 1 mM EDTA. Immunoprecipitation was performed using anti-FLAG M2 magnetic beads (Sigma) for a period of 3 hours with rotation at 4°C. Magnetic beads were isolated with a DynaMag-2 magnet (Thermo Fisher) and washed prior to addition of 1X NuPAGE LDS sample buffer and 1X NuPAGE Sample Reducing Reagent (Invitrogen). The non-immunoprecipitated whole cell lysates and immunoprecipitated fractions were heat denatured at 90°C for 15 minutes and 12 μL and 5 μL, respectively, were loaded into Criterion XT 12% Bis-Tris polyacrylamide gels (Bio-Rad) for SDS-PAGE using NuPAGE MES SDS Running Buffer (Invitrogen). Proteins were transferred Amersham Protran Premium Nitrocellulose Membrane, pore size 0.20 μm (GE Healthcare). Membranes were blocked with Odyssey blocking buffer in PBS (Li-COR) and incubated with anti-FLAG M2 (F1804; Sigma) and anti-c-Myc (C3956; Sigma). Secondary antibodies conjugated to DyLight 800 or 680 (Li-COR) and the Li-COR Odyssey imaging system were used to reveal specific protein detection. Images were analyzed and assembled using ImageStudioLite (Li-COR). To compare the steady-state expression levels of WT IFITM3 and mutants, HEK293T cells were transfected with 1.5 μg of empty pQCXIP or pQCXIP-FLAG-IFITM3 (encoding WT or the indicated mutants) and expression was measured using anti-FLAG M2 and analysis by flow cytometry or western blot analysis. For the former, cells were fixed/permeabilized with Cytofix/Cytoperm (BD) at 48 hours post-transfection, immunostained with anti-FLAG M2 (F1804; Sigma) and analyzed on a LSRFortessa (BD). For the latter, cells were lysed and separated by SDS-PAGE as indicated above and immunoblotting was performed with anti-FLAG M2 (F1804; Sigma) and anti-actin (C4, sc-47778; Santa Cruz Biotechnology). For blotting of HIV-1 proteins in cell lysates and concentrated virus supernatants, cell/virion lysis was performed with radioimmunoprecipitation (RIPA) buffer (Thermo Fisher) supplemented with Halt Protease Inhibitor mixture EDTA-free (Thermo Fisher) or 0.01% Triton-X (Sigma), respectively, and lysates were complemented with 1X NuPAGE LDS sample buffer and 1X NuPAGE Sample Reducing Reagent (Invitrogen) prior to heat denaturation at 90°C for 15 minutes. Samples were migrated and transferred as indicated above and immunoblotting was performed with the following antibodies: anti-p24 CA (NIH AIDS Reagent Resource: 3537), anti-gp120b (NIH AIDS Reagent Resource: 288), anti-gp41 2F5 (NIH AIDS Reagent Resource: 1475), anti-actin (C4, sc-47778; Santa Cruz Biotechnology), and anti-FLAG M2 (F1804; Sigma).

### Blue native PAGE

HEK293T cells were seeded in 6-well plates (750,000 per well) overnight and transfected with 1 μg empty pQCXIP or pQCXIP-FLAG-IFITM3 (either WT or G95L). At 48 hours post-transfection, cells were lysed with 1X NativePAGE Sample Buffer (Invitrogen) containing 1% digitonin. Lysates were mixed with 5% Coomassie G-250 at a volume to volume ratio of 20:1. Approximately 20 μg of protein per sample was loaded into a NativePAGE Novex 4-16% Bis-Tris polyacrylamide gel (Invitrogen) according to manufacturer’s instructions. A NativeMark Unstained Protein Standard was loaded as reference ladder. Following PAGE, proteins were transferred to Immobile FL PVDF membrane (EMD Millipore) and immunoblotting was performed with anti-FLAG M2 (F1804; Sigma). A secondary antibody conjugated to DyLight 800 or 680 (Li-COR) and the Li-COR Odyssey imaging system were used to reveal specific protein detection. Images were analyzed and assembled using ImageStudioLite (Li-COR).

### FRET and FLIM for oligomerization studies

HEK293T cells were seeded in μ-slide 8 well chambers (Ibidi) (50,000 per well) overnight and transiently co-transfected with 0.25 μg IFITM3-YFP and 0.25 μg mCherry or 0.25 μg IFITM3-YFP and 0.25 μg IFITM3-mCherry using TransIT-293 (Mirus). Constructs encoded WT IFITM3 or IFITM3 G95L and YFP/mCherry was fused to the amino-terminus. In parallel experiments, pairs of plasmids encoding PRRT2-YFP and PRRT2-mCherry were co-transfected. Constructs encoded WT PRRT2 or PRRT2 G305W and YFP/mCherry was fused to the carboxy-terminus. Living cells in Fluorobrite DMEM (Gibco) were imaged with a Zeiss LSM 780 confocal microscope using a 63X objective and oil immersion. To assess FRET, donor YFP fluorescence was detected with a gallium arsenide phosphide photomultiplier tube (GaAsP PMT) with a 520-550 nm emission window following excitation by a 514 nm laser. Acceptor mCherry fluorescence was detected with a GaAsP PMT detector with a 570-615 nm emission window following excitation by a 561 nm laser. FRET signal (acceptor mCherry fluorescence triggered by excitation of donor YFP) was collected with a GaAsP PMT detector with 570-615 nm emission window after excitation with a 514 nm laser. At least 50 cells per condition were examined in each experiment. Each cell was assigned a FRET index calculated using the FRET and colocalization analyzer plugin for Fiji (ImageJ). FLIM analysis of donor YFP was performed by excitation with a 950 nm two-photon, pulsed laser (Coherent) tuned at 80 MHz with single photon counting electronics (Becker Hickl) and detection with a HPM-100-40 module GaAsP hybrid PMT (Becker Hickl). Analysis was limited to cells exhibiting 250-1000 photons per pixel to mitigate the effects of photobleaching and low signal to noise ratio. SPCImage NG software (Becker Hickl) was used to acquire the fluorescence decay of each pixel, which was deconvoluted with the instrument response function and fitted to a Marquandt nonlinear least-square algorithm with two exponential models. The mean fluorescence lifetime was calculated as previously described [82] using SPCImage NG. At least 30 cells per condition were analyzed in each experiment.

### Number and Brightness analysis

HEK293T cells were seeded in μ-slide 8 well chambers (Ibidi) (50,000 per well) overnight and transiently with 0.50 μg IFITM3-mCherry using TransIT-293 (Mirus). Living cells in Fluorobrite DMEM (Gibco) were imaged with a Zeiss LSM 780 confocal microscope using a 63X objective and oil immersion. Regions of interest were limited to portions of cells which were immobile and which focused on plasma membrane fluorescence (intracellular signal from vesicular membranes was excluded). The axial position of a specimen during acquisition was stabilized using the Adaptive Focus Control module. mCherry was excited with a 561 nm laser and detected with a 570-615 emission window. For each cell, 100 frames were acquired at a frame rate of 0.385 frames per second with a 9.75 μs pixel dwell time and pixel size of 151.38 nm. Images were always acquired at 256 x 256 pixels such that the pixel size remained 3-4 times smaller than the volume of the point-spread function. Photobleaching of fluorescent proteins during data acquisition was corrected using a detrending algorithm [64]. Twenty cellular regions were examined per condition. Pixel-by-pixel brightness values were calculated in Fiji (ImageJ).

### FLIM for study of membrane order with FliptR

HEK293T stably expressing empty pQCXIP or pQCXIP-WT IFITM3 or the indicated mutant were seeded in μ-slide 8 well chambers (Ibidi) (50,000 per well) overnight and stained with 1 μM FliptR (Spirochrome) for 5 mins according to the manufacturer’s protocol. Imaging was performed with a 63X objective under oil immersion on a Leica SP8-X-SMD confocal microscope. When indicated, cells were treated with 100 μg/mL soluble cholesterol (C4951, Sigma), 5 mM methyl-cyclo-beta-dextrin (MßCD) (C4555, Sigma), or 1 μM Amphotericin B (A2942, Sigma) for one hour prior to addition of FliptR and imaging. Fluorescence was detected by hybrid external detectors in photon counting mode following excitation by a 488 nm pulsed laser turned to 20 MHz with single photon counting electronics (PicoHarp 300). Analysis was limited to cells with at least 250-1000 photons per pixel to mitigate the effects of photobleaching and low signal to noise ratio. Fluorescence decay of each pixel in FliptR-stained cells was acquired by Symphotime 64 software (Picoquant), and deconvoluted with the instrument response function and fitted to a Marquandt nonlinear least-square algorithm with two exponential models. The mean fluorescence lifetime (τ), in addition to individual component lifetimes (long and short lifetimes, τ_1_ and τ_2_), were calculated using Symphotime. At least 30 cells per condition were analyzed in each experiment.

## Figures and Figure Legends

**Supplemental Figure 1:**
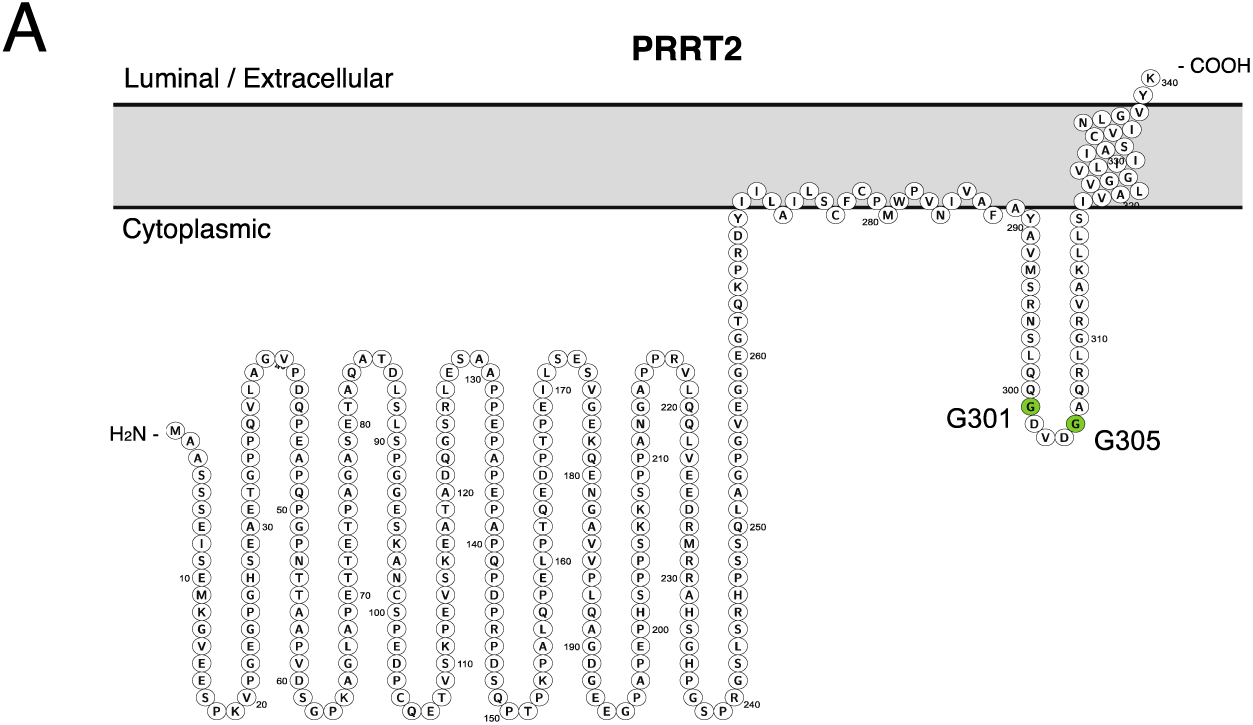
Membrane topology of PRRT2. (A) Schematic representation of the membrane topology of IFITM3 made with Protter. Residues corresponding to the glycines of GxxxG motif are indicated in green.

**Supplemental Figure 2:**
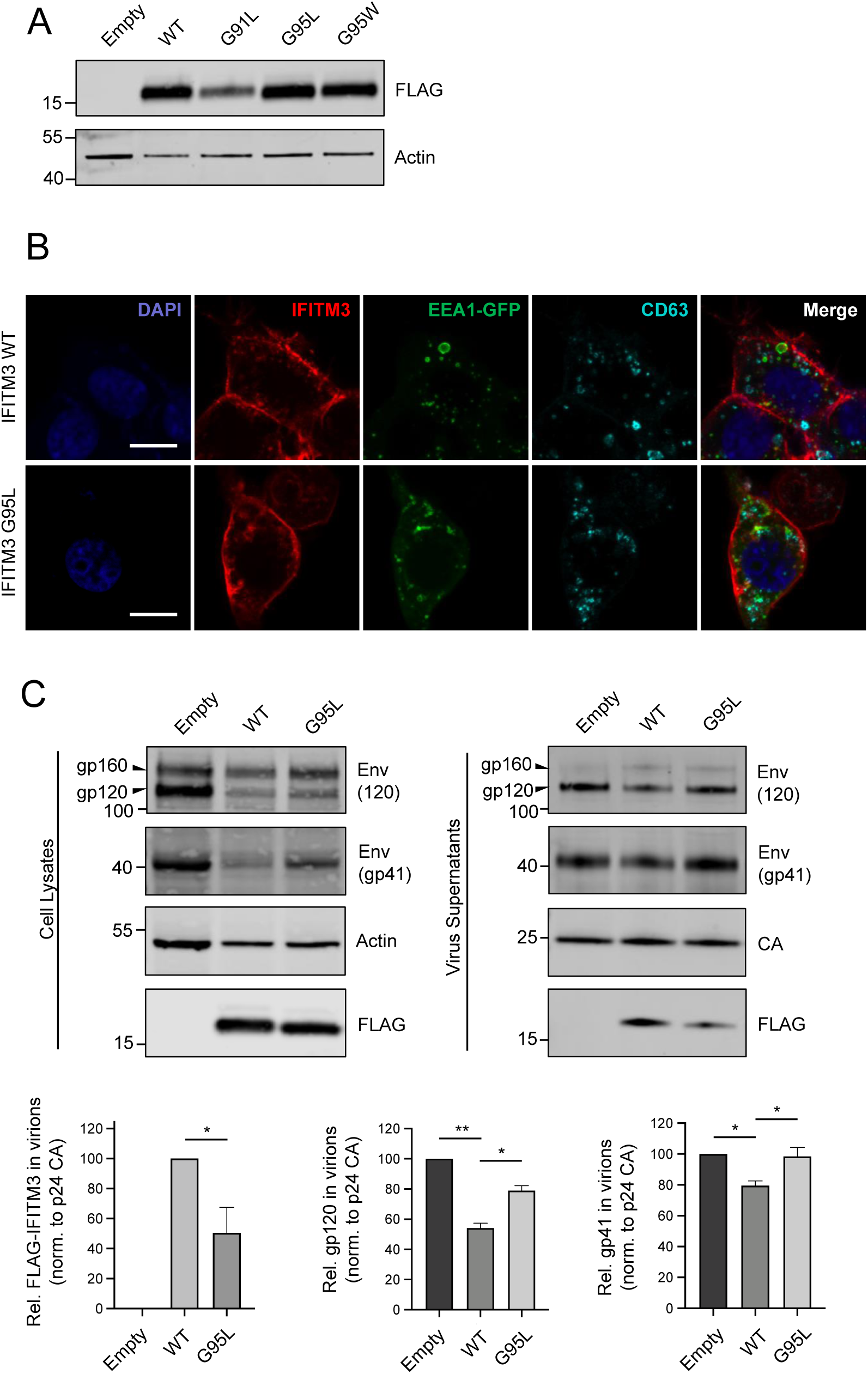
Expression, subcellular localization, and anti-HIV activities of WT and mutant IFITM3. (A) SDS-PAGE and Western blot analysis of whole cell lysates produced from HEK293T transiently transfected with empty pQCXIP, WT IFITM3-FLAG, or mutants. Immunoblotting was performed with anti-FLAG. Actin served as a loading control. Number and tick marks indicate size (kilodaltons) and position of protein standards in ladder. (B) HEK293T cells were co-transfected with EEA1-GFP and empty pQCXIP or IFITM3-FLAG (encoding WT or G95L). Cells were fixed, permeabilized, and immunostained at 48 hours post-transfection with anti-IFITM3 and anti-CD63 and cells were analyzed by immunofluorescence confocal microscopy. All images are average Z-stacks from 10 consecutive medial sections. Scale bars, 10 μm. (C) Whole cell lysates and virus-containing supernatants were collected from HEK293T co-transfected with the HIV-1 molecular clone NL4.3 and empty pQCXIP, WT IFITM3-FLAG, or the indicated mutant at 48 hours post-transfection. Virus-containing supernatants were ultracentrifuged through sucrose cushions. Both lysates and concentrated, purified virus-containing supernatants were subjected to SDS-PAGE and Western blot analysis. Immunoblotting was performed with anti-gp120, anti-gp41, anti-CA, anti-actin, and anti-FLAG. Levels of IFITM3 (FLAG), gp120, and gp41 were quantified and normalized to levels of CA. For anti-FLAG immunoblotting, the amount of WT IFITM3 in virions was set to 100%. For anti-gp120 and anti-gp41, levels observed in the pQCXIP empty vector were set to 100%.

**Supplemental Figure 3:**
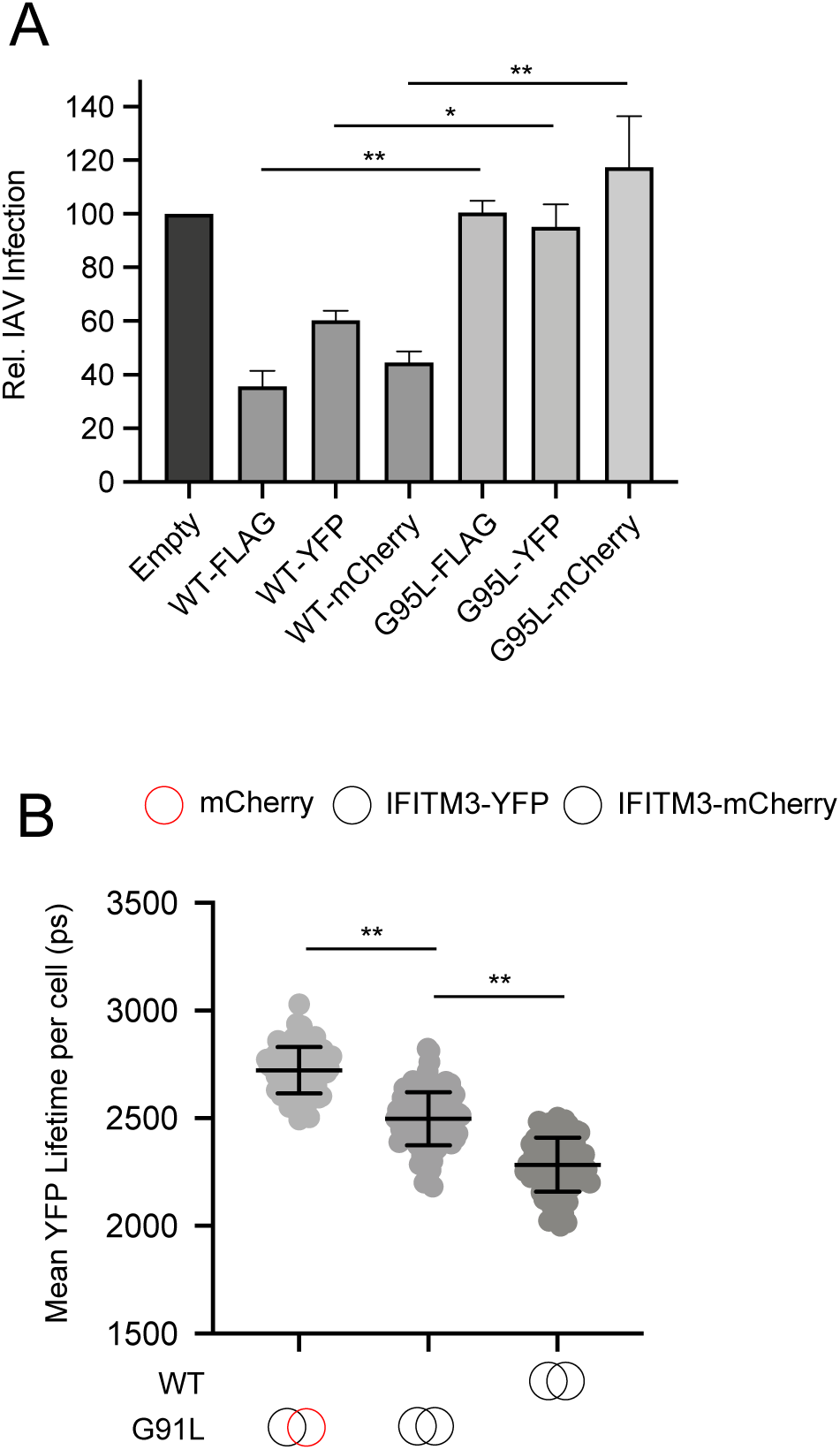
Antiviral activity of differentially-tagged IFITM3 and effect of glycine-91 on IFITM3 oligomerization. (A) HEK293T were transiently transfected with 1.5 micrograms of empty pQCXIP, FLAG-, YFP-, or mCherry-tagged IFITM3 (either WT or G95L) and, 48 hours later, challenged with IAV PR8 strain at a MOI of 0.1. Cells were fixed, stained with an anti-nucleoprotein antibody, and assessed by flow cytometry at 18 hours post-infection. Results represent the mean of three independent experiments. Error bars indicate standard error. (B) HEK293T were transiently co-transfected with IFITM3-YFP and mCherry or IFITM3-YFP and IFITM3-mCherry. Constructs encoded WT IFITM3 or IFITM3 G95L. Whole-cell YFP lifetimes were measured on a minimum of 50 cells per condition and the results of three independent experiments were pooled. Dots correspond to individual cells. Empty red circles are used to depict mCherry, filled red circles are used to depict IFITM3-mCherry (either WT or G95L), and filled yellow circles are used to depict IFITM3-YFP (either WT or G95L). Statistical analysis was performed using one-way ANOVA. *, P < 0.05; **, P < 0.001.

**Supplemental Figure 4:**
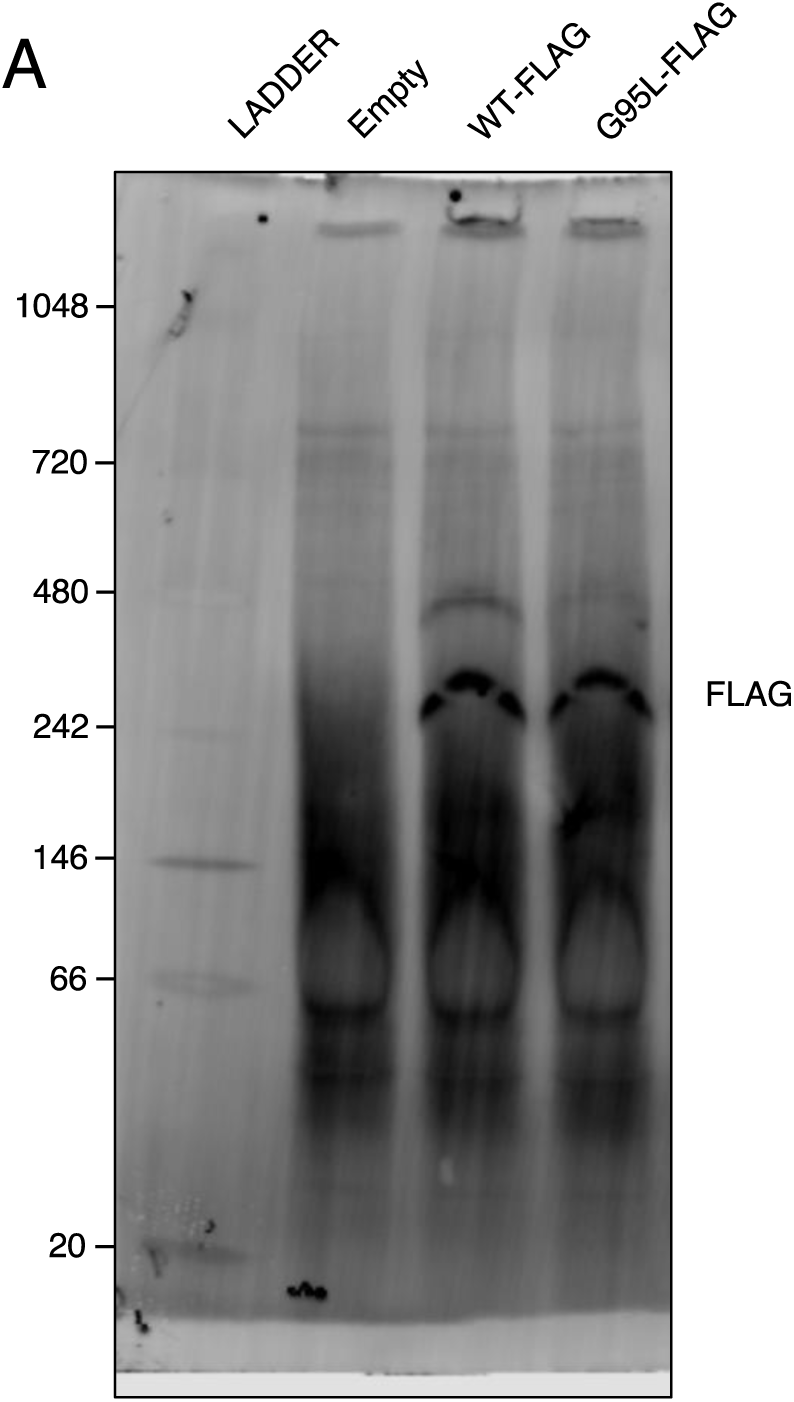
Blue native PAGE of IFITM3. (A) A full scan of the membrane depicted in Figure 4C is shown. Number and tick marks indicate size (kilodaltons) and position of protein standards in ladder.

**Supplemental Figure 5:**
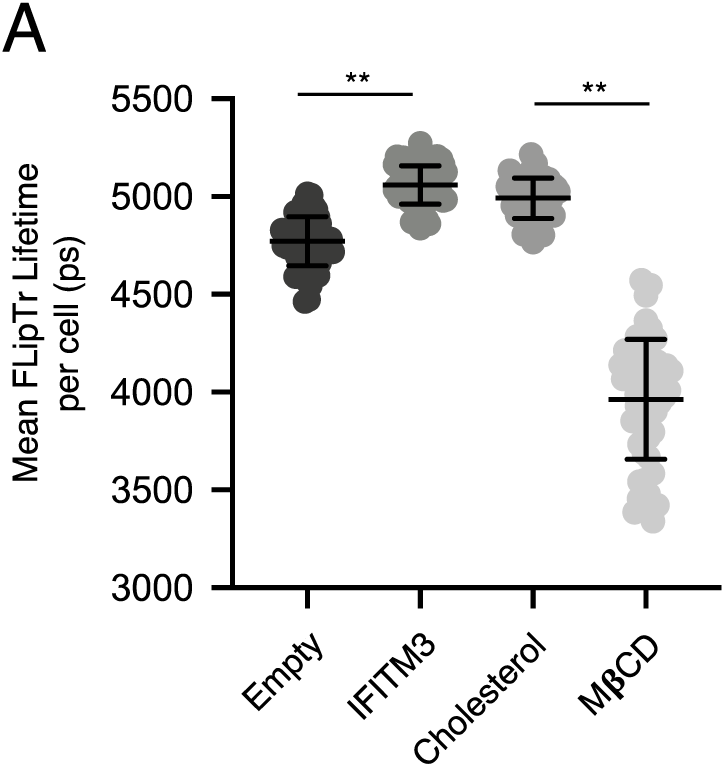
Assessment of cholesterol addition and cholesterol depletion on membrane order. (A) As in Figure 5A, except that soluble cholesterol (100 μg/mL) or methyl-beta-cyclo-dextrin (5 mM) were added to untransfected HEK293T cells for one hour and washed away prior to addition of 1 μM FliptR and cells were imaged by FLIM. τ_1_ was measured for a minimum of 40 cells per condition and the results of three independent experiments were pooled. Dots correspond to individual cells. Error bars indicate standard deviation. Statistical analysis was performed using one-way ANOVA. *, P < 0.05; **, P < 0.001.

## Acknowledgements

We thank Stephen Lockett and the Optical Microscopy and Image Analysis Laboratory of the National Cancer Institute, Center for Cancer Research, for technical support.

